# Targeting HIV at its core: A mathematical model for optimizing Tat Inhibitor-based therapies toward enhanced functional cure strategies

**DOI:** 10.64898/2026.04.13.718184

**Authors:** Rachel Waema, Cecilia Adongo, Sally Lago, Kennedy Ogutu

## Abstract

Human immunodeficiency virus (HIV) persistence remains a major barrier to cure due to the existence of long-lived latent reservoirs that evade immune clearance and persist despite combination antiretroviral therapy (ART). Although ART effectively suppresses viral replication, treatment interruption often leads to rapid viral rebound originating from these latent reservoirs. In this study, we develop a deterministic mathematical model describing the *in vivo* dynamics of HIV infection incorporating uninfected CD4^+^ T cells, infected cells, latent reservoirs, deep latent reservoirs, and infectious and non-infectious virions, while explicitly accounting for the therapeutic effects of reverse transcriptase inhibitors (RTIs), protease inhibitors (PIs), and Tat transcription inhibitors.

Analytical results establish positivity and boundedness of solutions and derive the effective reproduction number *R*_*e*_ using the next-generation matrix approach. Stability analysis shows that the virus-free equilibrium is locally asymptotically stable when *R*_*e*_ < 1, while viral persistence occurs when *R*_*e*_ > 1. Numerical simulations were performed to investigate therapy interactions, viral rebound following treatment interruption, and the impact of drug efficacy on viral set-points and latent reservoir dynamics.

To further explore therapy interactions, three-dimensional viral set-point surfaces and heat maps were generated to examine how combinations of infection inhibition, viral production inhibition, and transcriptional inhibition influence viral dynamics. The simulations reveal that Tat inhibition suppresses viral transcription, thereby reducing the transition of infected cells into productive infection and limiting viral propagation when combined with conventional ART mechanisms. The therapy parameter planes further demonstrate that strong transcriptional inhibition promotes the transition of infected cells into deep latency, supporting the emerging block-and-lock strategy for functional HIV cure. In addition, a three-dimensional eradication boundary surface and therapy cube were constructed to identify regions of parameter space where *R*_*e*_ < 1, corresponding to successful viral control. These visualizations show that viral eradication is unlikely when therapies act independently but becomes achievable when multiple therapeutic mechanisms act simultaneously.

Overall, the results highlight the critical role of transcriptional inhibition through Tat-targeting therapies in complementing existing ART regimens. By simultaneously suppressing viral replication and promoting deep latency, Tat-based combination strategies may significantly reduce viral rebound and contribute to long-term functional control of HIV infection.

## 1 Introduction

Human immunodeficiency virus (HIV) continues to pose a global health challenge, with over 39 million people currently living with the virus worldwide [1]. Despite major advances in antiretroviral therapy (ART), Human Immunodeficiency Virus (HIV) infection remains incurable due to the persistence of long-lived latent viral reservoirs. Combination ART effectively suppresses active viral replication and reduces plasma viral loads to undetectable levels, enabling individuals living with HIV to achieve near-normal life expectancy. However, ART does not eliminate latently infected cells harboring transcriptionally silent proviruses, which can reactivate following treatment interruption and lead to rapid viral rebound [2, 3]. Consequently, the latent reservoir remains the primary barrier to achieving either a sterilizing cure or durable ART-free remission. Latent HIV reservoirs are primarily established in long-lived memory CD4^+^ T cells where proviral genomes persist in a transcriptionally silent state. These reservoirs are maintained through multiple mechanisms, including epigenetic silencing, restricted transcriptional elongation, and limited availability of viral transcriptional activators such as the Tat protein [3, 4]. Upon cessation of therapy, stochastic reactivation of these latent proviruses results in rapid viral rebound, often occurring within weeks of ART interruption [2]. This phenomenon highlights the complex interplay between viral transcriptional regulation, host cellular mechanisms, and therapeutic interventions. Recent research has focused on several complementary strategies aimed at eliminating or controlling the HIV reservoir. One prominent approach is the “shock and kill” strategy, which employs latency reversing agents (LRAs) to induce viral transcription in latently infected cells, followed by immune-mediated clearance or cytopathic effects. However, many currently available LRAs demonstrate limited potency or lack specificity for HIV, limiting their effectiveness in clinical settings [4]. Alternatively, the “block and lock” strategy seeks to permanently silence viral transcription by targeting regulatory mechanisms such as Tat-mediated transcriptional elongation, thereby preventing viral reactivation even after therapy interruption. By broadly targeting HIV transcription, this approach is expected to affect not only the replication-competent reservoir but also the transcription and translation-competent proviral reservoir [5]. Although the majority of proviruses present in patients are defective and incapable of producing infectious progeny virions, a substantial fraction can still express viral RNA and proteins [6]. This residual viral protein production contributes to chronic immune activation and inflammation, even under long-term suppressive antiretroviral therapy [7]. Therefore, durable silencing of both replication-competent and defective proviruses may reduce persistent immune activation, limit viral antigen exposure, and ultimately confer additional clinical benefits beyond viral suppression, thereby improving overall patient health and quality of life [6].

The HIV transactivator protein Tat plays a central role in viral transcriptional regulation by recruiting the host transcriptional elongation factor P-TEFb to the viral promoter, thereby overcoming transcriptional pausing and enabling efficient viral gene expression. Increasing evidence suggests that Tat availability is a key determinant of whether HIV remains latent or becomes transcriptionally active. Recent studies have demonstrated that manipulating Tat expression or function can significantly influence latency reversal and viral reactivation dynamics [4, 8]. For example, nanoparticle-based delivery of Tat RNA has been shown to potently reactivate latent HIV in primary CD4^+^ T cells, highlighting the potential of Tat-targeted interventions as part of HIV cure strategies [9].

Since its discovery, substantial progress has been made in understanding the molecular biology, pathogenesis, and epidemiology of HIV, as well as the host’s immune response to the virus [10]. HIV, a retrovirus (as shown in Fig. 1), primarily targets CD4^+^ T cells. It does so by binding its gp120 surface protein to the CD4 receptor, enabling entry into the host cell [11]. Once inside, the viral RNA is reverse-transcribed into DNA and integrated into the host genome, establishing a lifelong infection. The integrated provirus is then transcribed into RNA, exported to the cytoplasm, and translated into viral proteins. Immature virions are assembled and bud from the plasma membrane, after which HIV protease cleaves viral polyproteins to mediate virion maturation [1].

**Fig 1.**
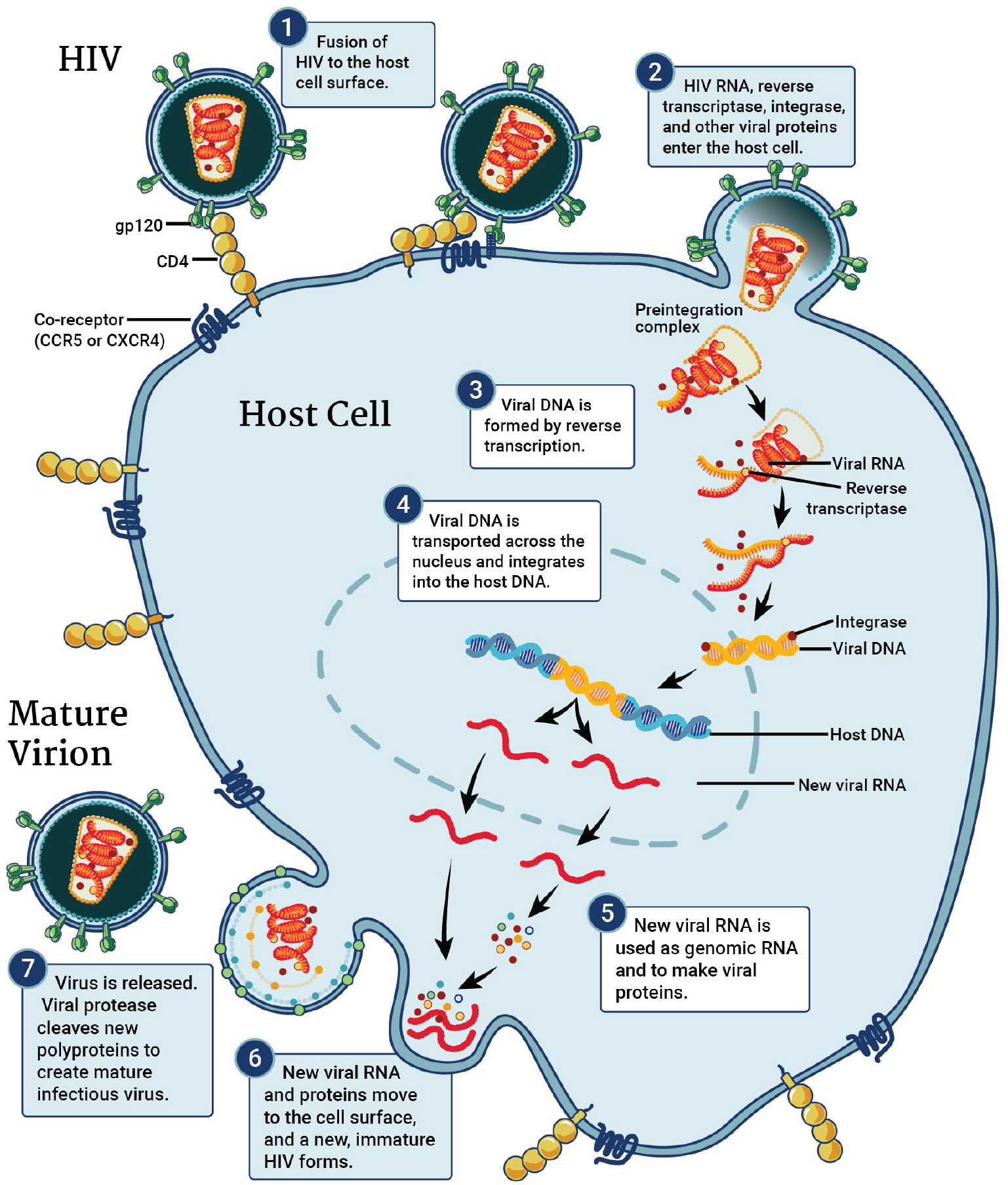
Stages of the HIV life cycle targeted by existing antiretroviral drugs. Reproduced from Chou TC, Maggirwar NS, Marsden MD. HIV Persistence, Latency, and Cure Approaches: Where Are We Now? Viruses. 2024;16:1163, licensed under the Creative Commons Attribution (CC BY 4.0) license.

In parallel with experimental advances, mathematical modeling has become an essential tool for understanding HIV persistence and evaluating potential cure strategies [12, 13]. Mathematical models allow researchers to investigate complex viral-host interactions that are difficult to study experimentally, particularly those involving latent reservoirs, treatment interruptions, and multi-drug therapeutic combinations. Recent modeling efforts have emphasized the importance of integrating viral transcriptional regulation, reservoir dynamics, and therapeutic efficacy in order to predict long-term treatment outcomes and identify optimal intervention strategies [14, 15, 16]. Such models can help quantify the conditions under which viral eradication becomes feasible or determine thresholds that prevent viral rebound after treatment interruption. A promising approach involves the use of Tat inhibitors, which target the HIV Tat protein, a critical regulator of viral transcription. By inhibiting Tat, these compounds can suppress viral replication and potentially limit the formation or maintenance of latent reservoirs [17]. However, the efficacy of Tat inhibitors and their integration into existing ART regimens remain active areas of investigation.

In this study, we develop and analyze a mathematical model describing HIV infection dynamics with explicit incorporation of latent reservoir compartments and therapeutic interventions targeting viral replication and transcriptional regulation. The model captures key biological processes including infection of susceptible cells, formation of latent reservoirs, viral production, and the effects of therapeutic agents that inhibit viral transcription and replication. Using this framework, we systematically investigate the impact of therapy combinations on viral persistence, reservoir dynamics, and potential eradication outcomes. By integrating mathematical modeling with biologically informed parameter exploration, this work provides new insights into the mechanistic drivers of viral persistence and the potential role of transcription-targeting therapies in HIV cure strategies. The insights gained from this model will inform clinical decision-making, help refine HIV treatment protocols and advance global efforts toward achieving a functional cure for HIV [15, 18].

## 2 Materials and Methods

To describe HIV/AIDS in-vivo progression throughout the viral life cycle and under various stages of transcription, as well as the relative impact of therapeutic interventions, the host’s immune system and the viral dynamics are divided into seven compartments representing different populations of CD4^+^ T cells and virion particles. The compartments represent uninfected CD4^+^ T cells *T*(*t*), infected CD4^+^ T cells *I*(*t*), latently infected CD4^+^ T cells *I*_*L*_(*t*), deeply latent CD4^+^ T cells *I*_*D*_(*t*), productively infected CD4^+^ T cells *I*_*P*_(*t*), infectious virion particles *V*_*I*_(*t*), and non-infectious virion particles *V*_*U*_(*t*). The model captures the interactions between CD4^+^ T cells and the HIV virus at various stages of infection, highlighting different latency stages and the influence of enzymes like reverse transcriptase, Trans-Activation protein (Tat), and protease. It also incorporates the impact of therapeutic interventions, including reverse transcriptase inhibitors, Tat inhibitors, and protease inhibitors.

The uninfected CD4^+^ T cell population *T*(*t*) begins with cells recruited from the thymus at a rate Π, and these cells undergo natural death at a rate *µ*_*T*_ . In this model, the recruitment of CD4^+^ T cells is restricted by a carrying capacity, reflecting the body’s limitations in replenishing CD4^+^ T cells. This carrying capacity is implemented by modifying the recruitment term Π as a function of the current CD4^+^ T cell population and the total carrying capacity *K*, such that Π 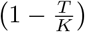 models the restricted recruitment when *T*(*t*) approaches *K*.

HIV virions infect uninfected CD4^+^ T cells at a rate *β*. Once infection occurs, infected cells *I*(*t*) may die naturally at a rate *µ*_*I*_, progress to productive infection through viral transcription at a rate (1 − *ω*)*α*, or enter latency at a rate *ρ*. The parameter *α* represents the viral transcription rate, while *ω* represents the effectiveness of the Tat inhibitor, which reduces transcription and limits the progression of infected cells to productive infection.

Latently infected CD4^+^ T cells *I*_*L*_(*t*) may either reactivate to productive infection at a rate (1 − *ω*)*γ*, transition into the deep latency state at a rate influenced by tat inhibitor *ω*, or die naturally at a rate *µ*_*L*_. The Tat inhibitor therefore limits reactivation while simultaneously promoting the transition of latent cells into the deeply latent compartment.

The deeply latent infected CD4^+^ T cells *I*_*D*_(*t*) represent transcriptionally silent cells that do not reactivate and can only be removed through natural death at a rate *µ*_*D*_. Productively infected cells *I*_*P*_(*t*) produce infectious and non-infectious virions at rates (1 − *E*)*κ* and *Eκ*, respectively, and die at a natural rate *µ*_*P*_ .

In summary, uninfected CD4^+^ T cells, infected CD4^+^ T cells, latently infected CD4^+^ T cells, deeply latent infected CD4^+^ T cells, and productively infected CD4^+^ T cells undergo natural death at their respective rates (*µ*_*T*_, *µ*_*I*_, *µ*_*L*_, *µ*_*D*_, and *µ*_*P*_). Both infectious and non-infectious virion particles are cleared from the body at the same rate *δ*. This assumes that the immune system and other mechanisms responsible for viral clearance do not differentiate between infectious and non-infectious particles.

This model not only describes the dynamics of HIV infection and latency but also accounts for the therapeutic effects of reverse transcriptase inhibitors, Tat inhibitors, and protease inhibitors. In particular, the model explicitly incorporates the role of Tat inhibition in reducing viral transcription and promoting the transition of latent infected cells into a deeply latent state, providing a detailed framework to evaluate how these interventions restrict viral replication and influence the persistence of latent reservoirs.

### Model Assumptions

To accurately describe the in-vivo progression of HIV/AIDS and assess the impact of various therapeutic interventions, the following assumptions are made in the mathematical model:

1. The virus is assumed to specifically target CD4^+^ T cells for infection, and other cell types are not considered as targets for HIV infection.
2. The infectious and non-infectious virus particles are being cleared at the same rate of *δ*.
3. The latent and deeply latent infected CD4^+^ T cells cannot produce new virions.
4. The efficacy of each therapeutic intervention (Reverse transcriptase inhibitors, Tat inhibitors and protease inhibitors) is assumed to remain constant throughout the course of treatment, without accounting for drug resistance or changes in the patient’s adherence to the treatment regimen.

The parameters and variables used in the model are described in Table 1 and 2 below.

**Table 1.** Variables for HIV in vivo model with therapy.

| Symbol | Description |
| --- | --- |
| $T$ | Concentration of healthy CD4<sup>+</sup> T cells per mm<sup>3</sup> at time $t$. |
| $I$ | Concentration of infected CD4<sup>+</sup> T cells at time $t$. |
| $I_L$ | Concentration of latently infected CD4<sup>+</sup> T cells per mm<sup>3</sup> at time $t$. |
| $I_P$ | Concentration of productively infected CD4<sup>+</sup> T cells at time $t$. |
| $I_D$ | Concentration of deeply latently infected CD4<sup>+</sup> T cells at time $t$. |
| $V_I$ | Concentration of infectious virions at time $t$. |
| $V_u$ | Concentration of non-infectious virions at time $t$. |

**Table 2.**
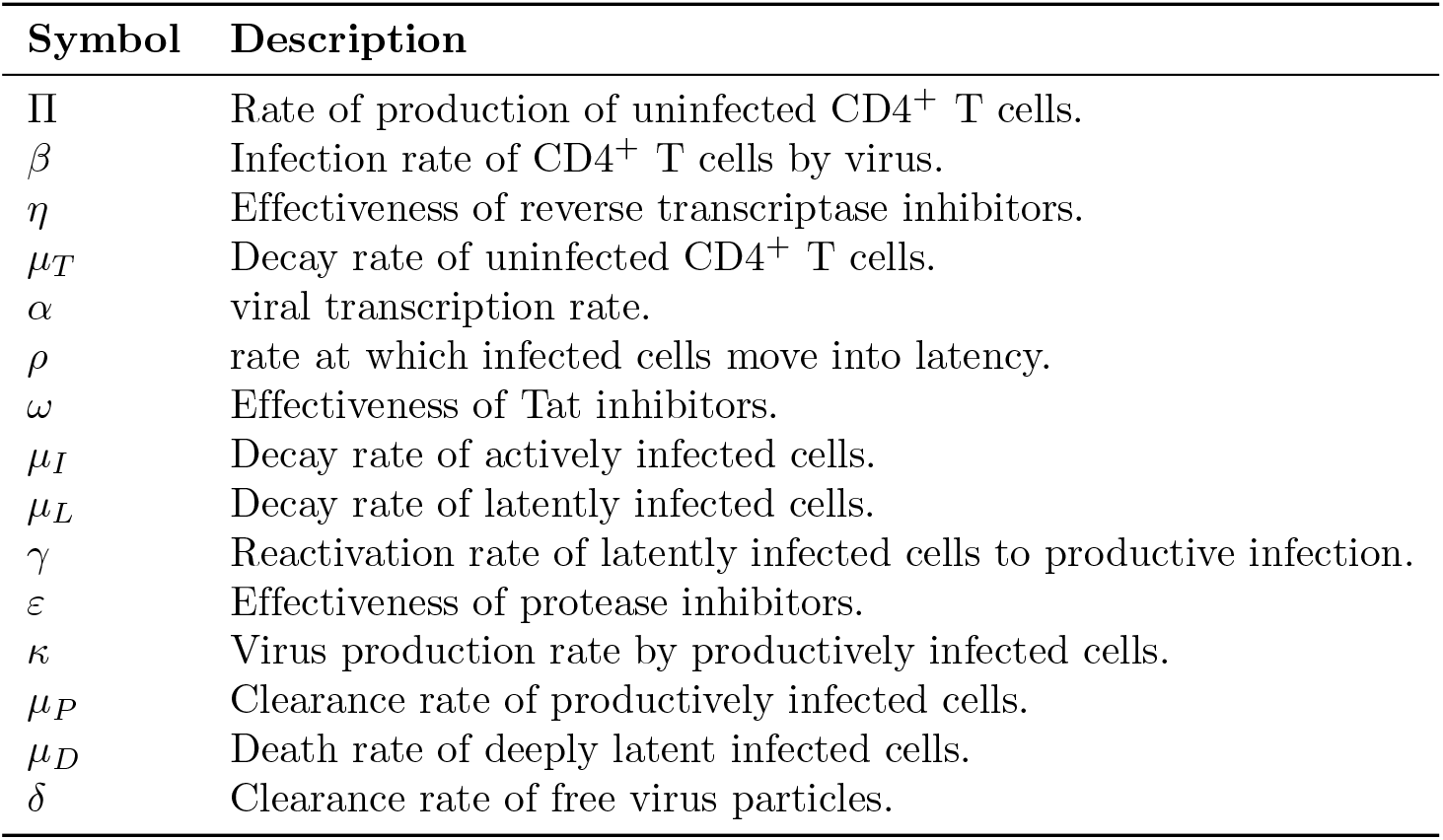
Parameters for HIV in vivo model with therapy.

The developed non-parameterized and parameterized models are illustrated in Figs 2 and 3 below.

**Fig 2.**
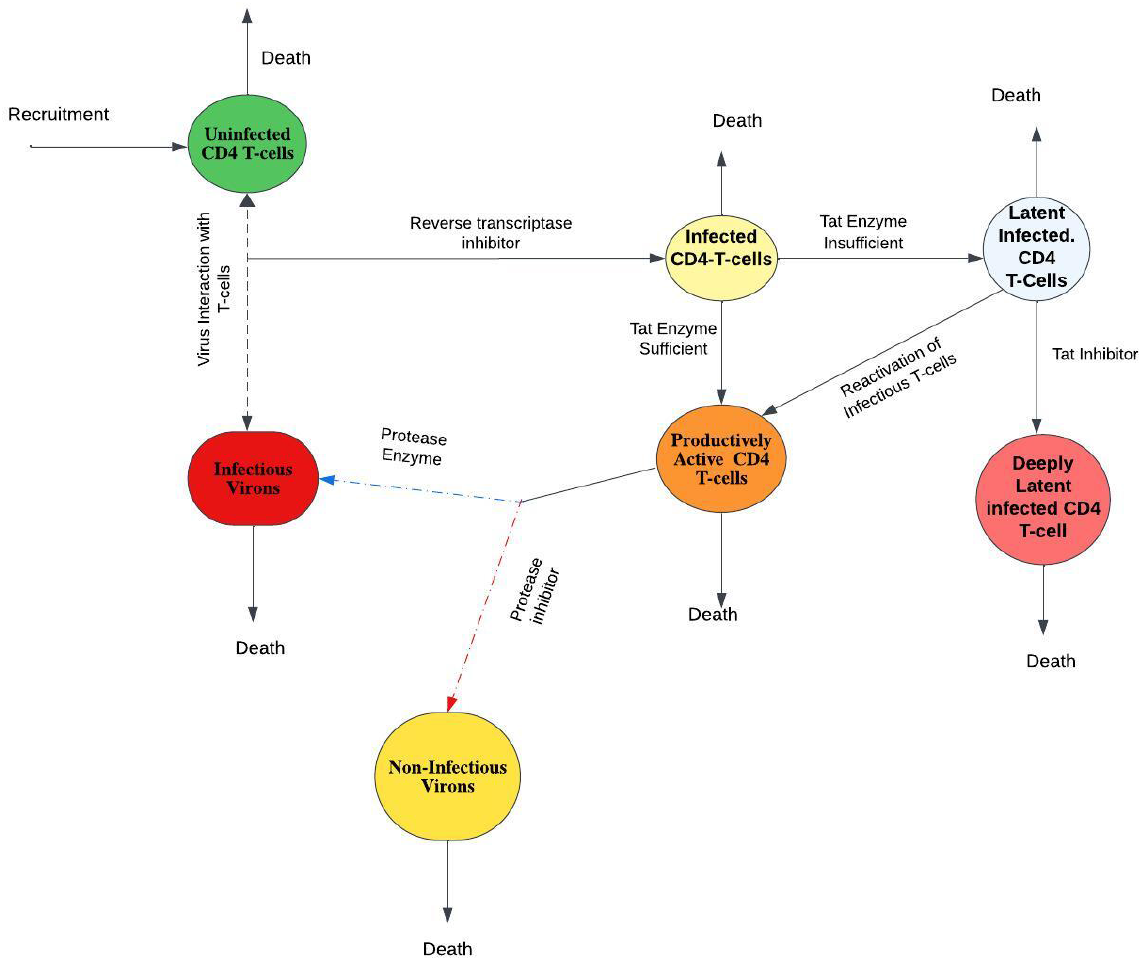
Non-Parameterized Model.

**Fig 3.**
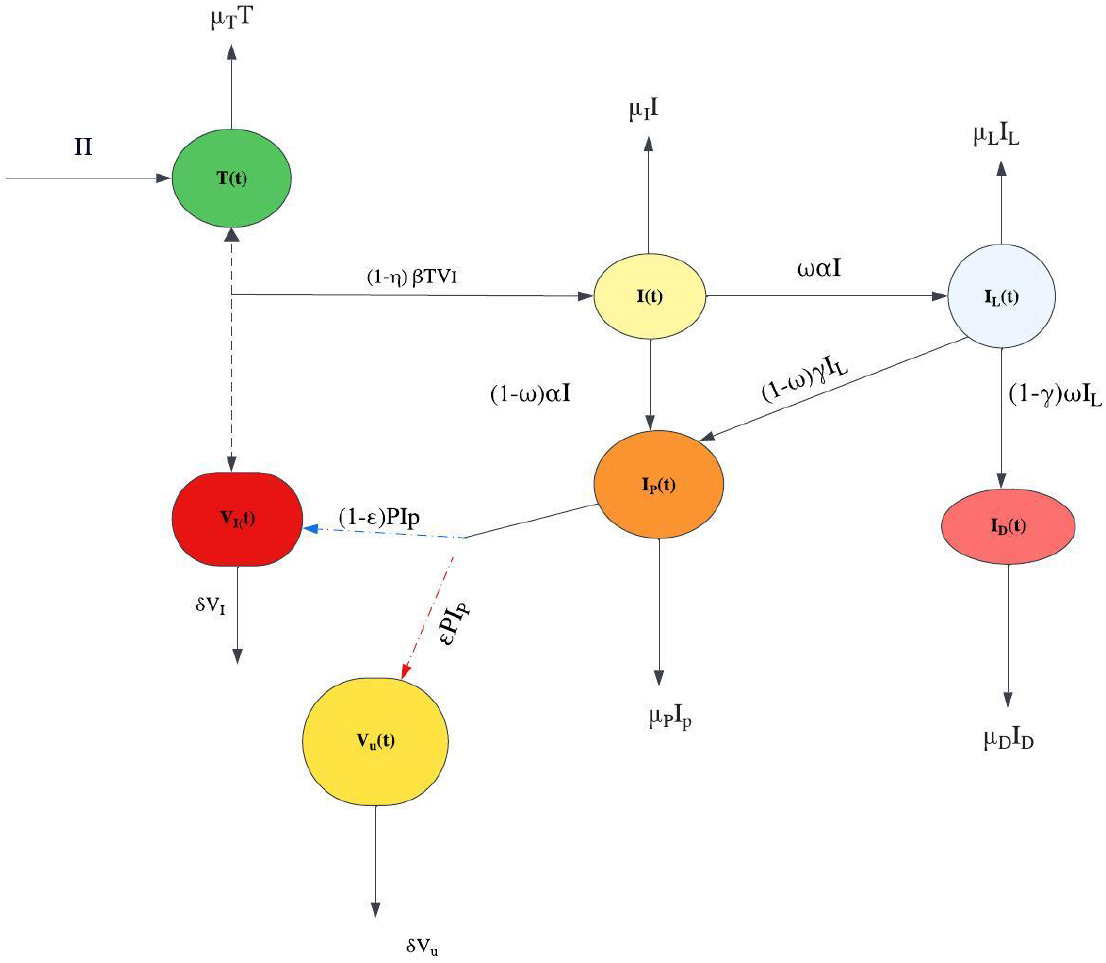
Parameterized Model.

The resulting non-linear system of differential equations are:

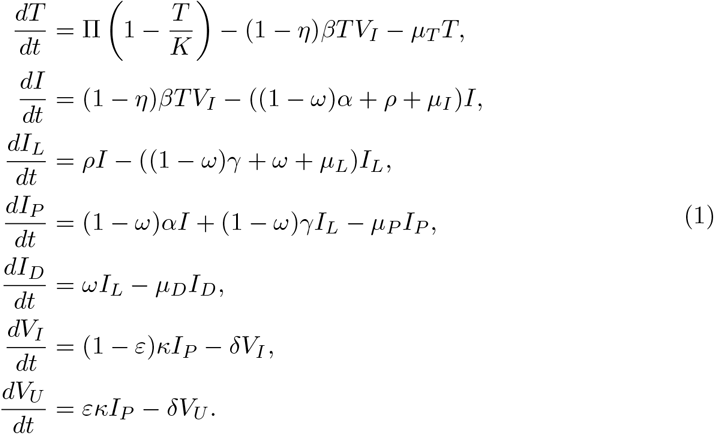

with initial conditions; *T*(0) = *T*_0_, *I*(0) = 0, *I*_*L*_(0) = 0, *I*_*P*_ (0) = 0,*I*_*D*_(0) = 0, *V*_*I*_ (0) = 0, *V*_*U*_ (0) = 0.

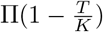 is the modified recruitment term for healthy CD4^+^ T cells, which incorporate the carrying capacity K. The carrying capacity K represents the maximum concentration of healthy CD4^+^ T cells that the body can sustain.

## 3 Model Analysis

This section covers the analysis of the model, focusing on key properties pertinent to HIV dynamics. Specifically, we have examined the positivity and boundedness of the solutions, ensuring their feasibility within the context of the model. To ensure biological feasibility, we first establish that all state variables of the model remain non-negative and bounded for all time.

### 3.1 Positivity of Solutions

The monitoring of cell populations in the in-vivo HIV model underscores the importance of verifying the non-negativity of its state variables. To clarify, we demonstrate that under initial conditions where all values are non-negative, the solutions to model (1) will remain non-negative for all times *t* ≥ 0. This leads us to the following theorem.

#### Theorem 1

*Let the parameters for model* (1) *be non-negative constants. A non-negative solution T*(*t*), *I*(*t*), *I*_*L*_(*t*), *I*_*D*_(*t*), *I*_*P*_(*t*), *V*_*I*_(*t*), *V*_*U*_(*t*) *for the model exists for all state variables with non-negative constants*

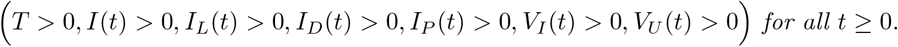

*Proof*. From the first equation of model (1)

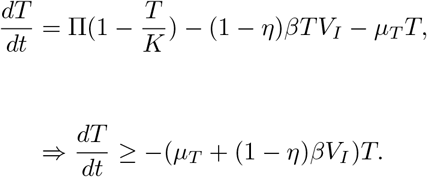

Therefore, by direct integration

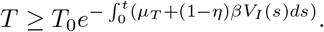

which implies, that

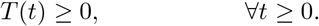

A similar comparison argument applies to the remaining state variables *I*(*t*), *I*_*L*_(*t*), *I*_*D*_(*t*), *I*_*P*_(*t*), *V*_*I*_(*t*), and *V*_*U*_(*t*), each of which can be bounded below by solutions of linear differential inequalities with non-negative initial data. Hence, all solutions of model (1) remain non-negative for all *t* ≥ 0 and for non–negative initial conditions.

### 3.2 Boundedness of Solutions

#### Theorem 2.

All solutions (*T*(*t*), *I*(*t*), *I*_*L*_(*t*), *I*_*D*_(*t*), *I*_*P*_(*t*), *V*_*I*_(*t*), *V*_*U*_(*t*)) ∈ R^7^ are bounded.

*Proof*. Let the total population of the CD4^+^ T cells be denoted by

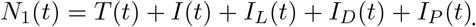

Summing the corresponding equations in system (1) yields

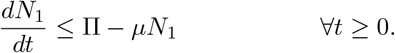

where *µ* = *min*(*µ*_*T*_, *µ*_*I*_, *µ*_*L*_, *µ*_*D*_, *µ*_*P*_). Solving this differential inequality shows that

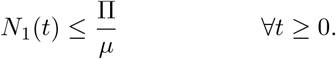

establishing uniform boundedness of the CD4^+^ T cells populations.

*Proof*. Similarly, let the total population of the virus be represented by

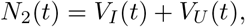

From system (1)

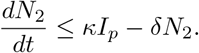

Since *I*_*P*_(*t*) is bounded by the bound on *N*_1_(*t*), it follows that

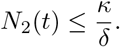

which can be re-written as

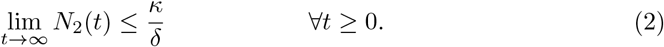

Therefore all state variables remain bounded in a biologically feasible region Ω = *{*(*T*(*t*), *I*(*t*), *I*_*L*_(*t*), *I*_*D*_(*t*), *I*_*P*_(*t*), *V*_*I*_(*t*), *V*_*U*_(*t*)) ≥ 0*}*

### 3.3 Virus Free Equilibrium and Effective Reproduction Number

#### 3.3.1 Virus-free Equilibrium

The virus-free equilibrium corresponds to the absence of infection in the host and is obtained by setting all infected cells and virus compartments to zero. Thus, setting

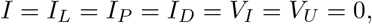

the virus-free equilibrium of system (1) is given by

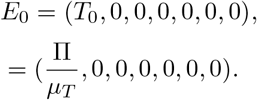

#### 3.3.2 Effective Reproduction Number

The effective reproduction number *R*_*e*_ is defined as the expected number of secondary infected cells produced by a single infected where interventions are in place. To derive *R*_*e*_, we employ the next-generation matrix method [19]. We consider all compartments that contribute to infection and persistence, namely:

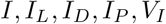

Let *F* denote the rate of appearance of new infections, *V* denote the rate of transfer of individuals between infected compartments and removal from infection. The infected subsystem of system (1) can be written as

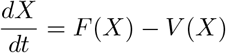

where *X* = (*I, I*_*L*_, *I*_*D*_, *I*_*P*_, *V*_*I*_)

#### 3.3.3 New Infection Terms

New infections arise solely through the infection of uninfected CD4^+^ T cells by infectious virions. Thus, the vector of new infection terms is given by

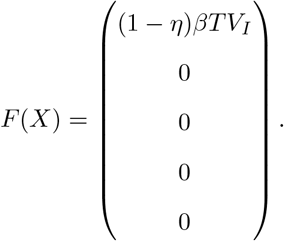

Evaluating the Jacobian of *F* at the virus-free equilibrium *E*_0_ yields

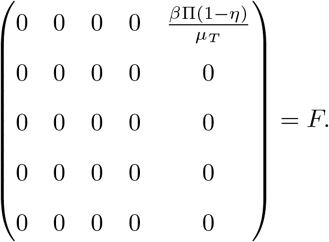

#### 3.3.4 Transition and Removal Terms

The remaining transition terms are given by:

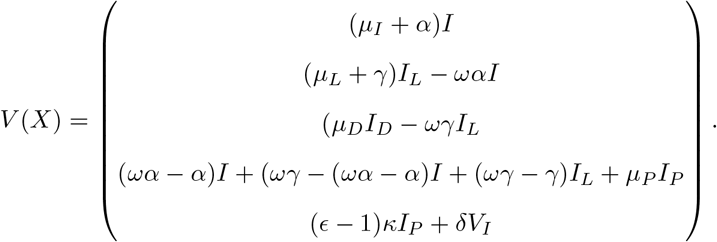

The Jacobian matrix of *V* evaluated at the virus-free equilibrium *E*_0_ is given by

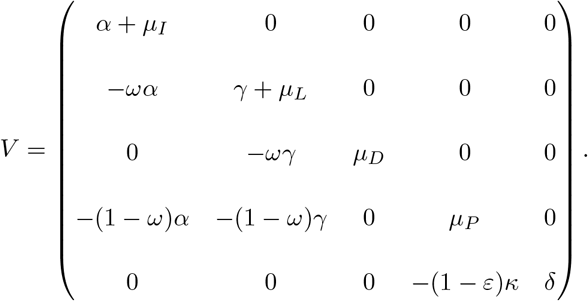

### 3.4 Computation of the Effective Reproduction Number

The next-generation matrix is defined as

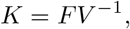

where *F* represents the matrix of new infection terms and *V* denotes the matrix of transition and removal terms. The effective reproduction number is given by the spectral radius of *K*,

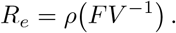

After algebraic simplification, the effective reproduction number is obtained as

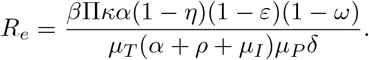

Which can be written as

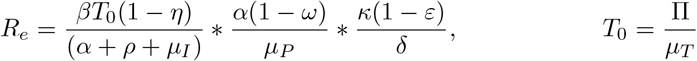

This expression shows that the persistence of HIV infection within the host is influenced by the effectiveness of the therapeutic interventions. An increase in the effectiveness of these interventions reduces the value of *R*_*e*_.

The parameter *η* represents the efficacy of reverse transcriptase inhibitors, which reduce the infection of CD4^+^ T cells by decreasing the effective infection rate. The parameter *ε* represents the efficacy of protease inhibitors, which reduce the production of infectious virions from productively infected cells. The parameter *ω* represents the effectiveness of Tat inhibitors, which suppress viral transcription and promote the transition of infected and latent cells toward deeper latency, thereby reducing progression to productive infection and leading to a decrease in the effective reproduction number *R*_*e*_.

If the combined therapeutic interventions reduce *R*_*e*_ below unity (*R*_*e*_ < 1), the infection cannot sustain itself and viral replication declines over time. However, if *R*_*e*_ > 1, the infection persists within the host despite treatment, highlighting the importance of combining multiple therapeutic strategies to effectively control HIV replication and reduce viral reservoirs.

### 3.5 Local Stability of the Disease-Free Equilibrium

The local stability of the virus-free equilibrium is investigated by linearizing system (1) about the virus-free equilibrium

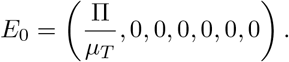

To determine the local stability of *E*_0_, we analyze the Jacobian matrix of the system evaluated at the virus-free equilibrium.

The dynamics of the infected compartments are governed by the effective reproduction number *R*_*e*_. Using the next generation matrix approach, the effective reproduction number is given by

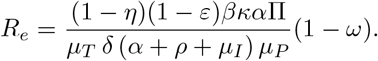

**Theorem**.

The virus-free equilibrium *E*_0_ is locally asymptotically stable if *R*_*e*_ < 1 and unstable if *R*_*e*_ > 1.

**Proof**.

The Jacobian matrix evaluated at *E*_0_ yields eigenvalues associated with the infected subsystem whose dominant eigenvalue is governed by *R*_*e*_. If *R*_*e*_ < 1, all eigenvalues have negative real parts, implying that small perturbations around the disease-free equilibrium decay over time. Consequently, the infection dies out and the system returns to *E*_0_. If *R*_*e*_ > 1, at least one eigenvalue has a positive real part, leading to growth of the infected compartments and persistence of infection in the host. Hence, the virus-free equilibrium is locally asymptotically stable whenever *R*_*e*_ < 1 and unstable whenever *R*_*e*_ > 1.

### 3.6 Global Stability of the Virus-Free Equilibrium

To establish the global stability of the virus-free equilibrium, we construct a suitable Lyapunov function.

Recall that the virus-free equilibrium of the system is given by

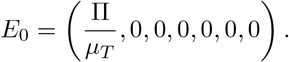

Consider the Lyapunov function defined by

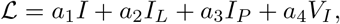

where *a*_1_, *a*_2_, *a*_3_, and *a*_4_ are positive constants to be determined. Taking the time derivative of ℒ along solutions of the system gives

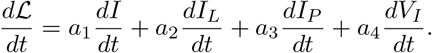

Substituting the model equations yields

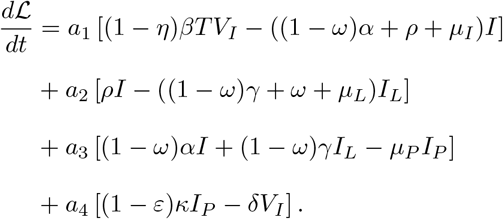

Evaluating the derivative near the virus-free equilibrium where

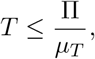

we obtain

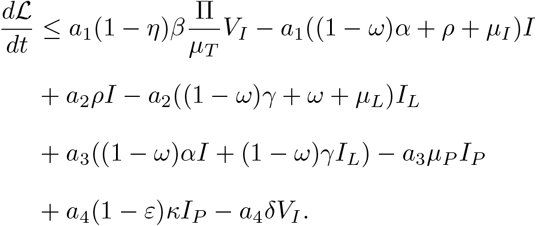

Choosing appropriate constants *a*_1_, *a*_2_, *a*_3_, *a*_4_ allows the positive terms to cancel and ensures

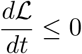

Whenever

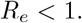

Thus,

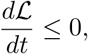

with equality holding only when

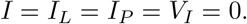

Therefore, the largest invariant set contained in

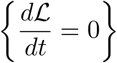

is the virus-free equilibrium *E*_0_.

By LaSalle’s Invariance Principle, all trajectories in the feasible region converge to *E*_0_ whenever *R*_*e*_ < 1.

**Theorem**.

If *R*_*e*_ < 1, the virus-free equilibrium *E*_0_ is globally asymptotically stable in the feasible region Ω.

The same applies when global stability approach of Castillo-Chavez and Song [20] for epidemic models is applied.

This indicates that sufficiently effective treatment strategies can suppress viral replication and eliminate active infection dynamics.

### 3.7 Sensitivity Analysis of *R*_*e*_

To understand the impact of model parameters on the potential spread of HIV within the host, we performed sensitivity analysis of the effective reproduction number, *R*_*e*_. We employed the Partial Rank Correlation Coefficient (PRCC) method to quantify the influence of each parameter on *R*_*e*_. PRCC is particularly suitable for nonlinear models and allows for the identification of parameters that strongly affect model outcomes while accounting for correlations among parameters.

Parameter ranges for the analysis were based on biological plausibility and literature values as shown in Table 3. The PRCC values were then computed between each parameter and *R*_*e*_, providing a ranked measure of sensitivity as shown in Fig 4. Parameters with larger absolute PRCC values indicate a stronger influence on the effective reproduction number.

**Table 3.** Parameter values for HIV in vivo model with therapy.

| Parameter | Value | Source |
| --- | --- | --- |
| $\Pi$ | 20 cells/mm<sup>3</sup>/day | [21] |
| $\beta$ | $2.4 \times 10^{-5}$ mm<sup>3</sup> vir<sup>-1</sup> day<sup>-1</sup> | [18] |
| $\mu_T$ | 0.01 day<sup>-1</sup> | [22] |
| $\alpha$ | 0.5 day<sup>-1</sup> | [23] |
| $\mu_I$ | 0.02 day<sup>-1</sup> | [24] |
| $\mu_L$ | 0.06 day<sup>-1</sup> | [25] |
| $\gamma$ | 0.05 day<sup>-1</sup> | Assumed |
| $\kappa$ | 100 vir cell<sup>-1</sup> day<sup>-1</sup> | [26] |
| $\rho$ | 0.5 day<sup>-1</sup> | [23] |
| $\mu_P$ | 0.05 day<sup>-1</sup> | [27] |
| $\mu_D$ | 0.07 day<sup>-1</sup> | Assumed |
| $\delta$ | 0.05 day<sup>-1</sup> | Assumed |
| $K$ | $1.0 \times 10^5$ cells/mm<sup>3</sup> | [28] |

**Fig 4.**
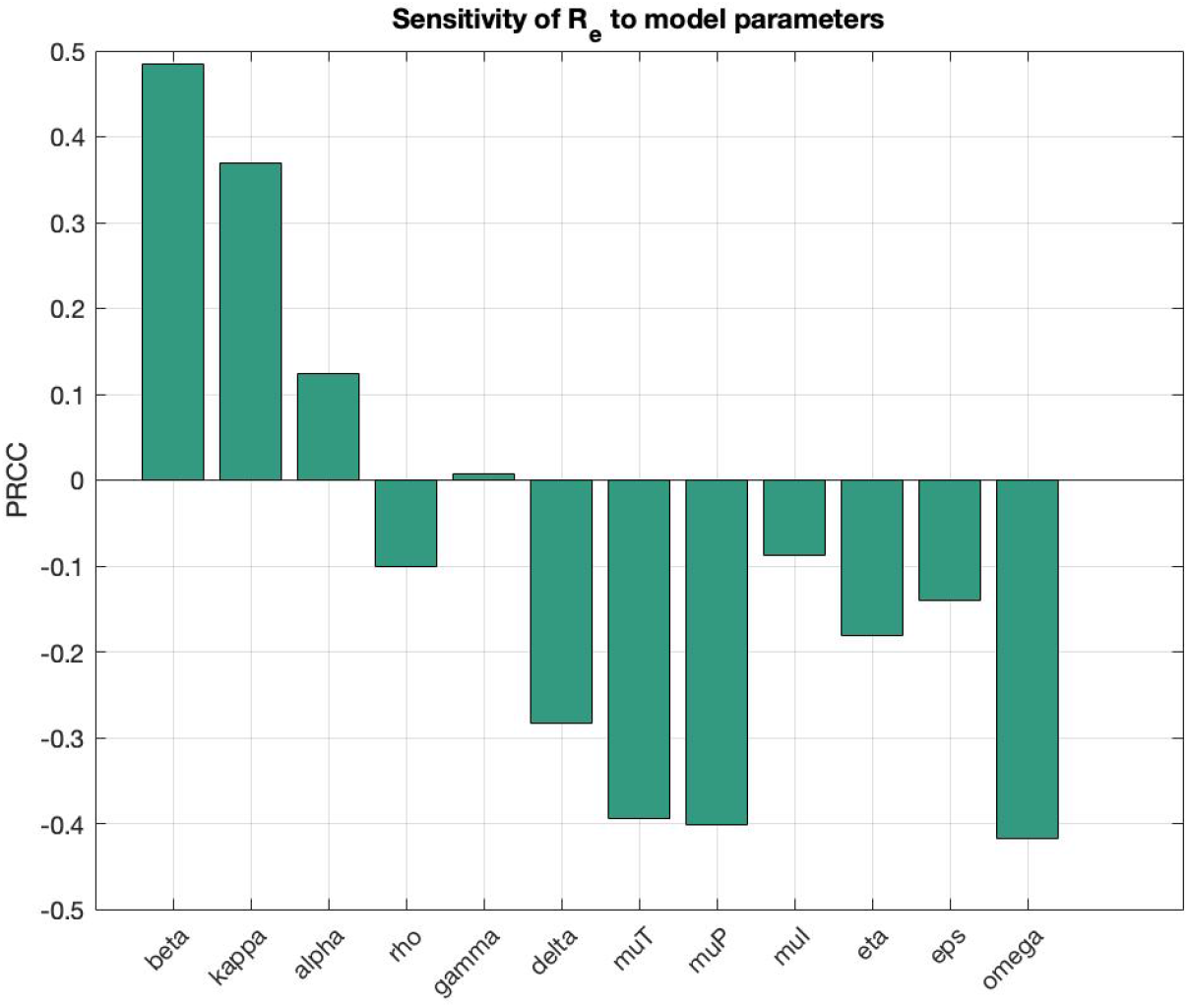
Partial Rank Correlation Coefficients (PRCC) of model parameters on the effective reproduction number *R*_*e*_. Positive values indicate an increase in *R*_*e*_ with increasing parameter value, while negative values indicate a decrease.

The analysis in Fig 4 allowed us to identify the most influential parameters controlling HIV dynamics in the model. From a biological perspective, this suggests that targeting processes related to viral production (*κ*), HIV infection rates (*β*), viral reactivation (*ω*) or enhancing mechanisms associated with virus or infected cell clearance (*δ, µ*_*P*_) could have the greatest impact on controlling infection dynamics.

### 3.8 Endemic Equilibrium and Bifurcation Analysis

In this section, we examine the existence and stability of the endemic (chronic infection) equilibrium of system (1) and describe the bifurcation structure governing the transition between viral clearance and persistent infection. This analysis provides further insight into the long-term within-host dynamics of HIV under different therapeutic interventions.

#### 3.8.1 Endemic Equilibrium

The endemic equilibrium *E*^∗^ corresponds to the steady state where infection persists in the host. It is obtained by setting the right-hand side of the system equal to zero. Let

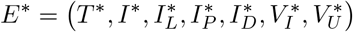

denote the endemic equilibrium of system (1). This equilibrium represents chronic HIV infection under sustained viral replication and latency dynamics. Setting the right-hand side of system (1) equal to zero, we obtain the equilibrium conditions. For *R*_*e*_ > 1, these conditions admit a unique endemic equilibrium *E*^∗^ which satisfies:

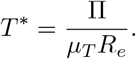

The remaining state variables can be written as

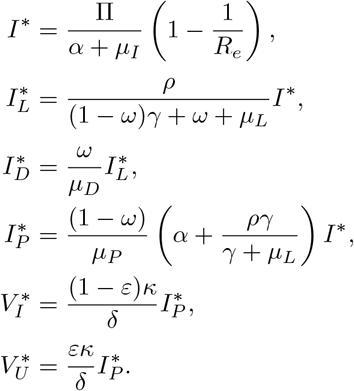

All components of *E*^∗^ are strictly positive whenever *R*_*e*_ > 1, and vanish continuously as *R*_*e*_ → 1, consistent with the forward bifurcation established below. The endemic equilibrium therefore exists whenever *R*_*e*_ > 1.

Expressing the endemic equilibrium in terms of *R*_*e*_ highlights the threshold structure of the model, that is therapeutic interventions reduce equilibrium viral burden primarily by driving *R*_*e*_ toward unity.

### 3.9 Stability of the Endemic Equilibrium

The stability of the endemic equilibrium is determined by analyzing the Jacobian matrix evaluated at *E*^∗^. Standard results in epidemiological modelling show that when *R*_*e*_ > 1, the endemic equilibrium exists and is locally asymptotically stable.

Thus, if *R*_*e*_ > 1, the virus persists within the host and the system approaches the endemic steady state.

Conversely, when *R*_*e*_ < 1, the endemic equilibrium does not exist and the infection is cleared.

#### 3.9.1 Bifurcation Analysis at the Threshold *R*_*e*_ = 1

To further understand the impact of therapeutic interventions on HIV dynamics, we perform a bifurcation analysis of system (1) with respect to key parameters influencing viral replication and latency formation. In particular, we focus on the Tat inhibitor effectiveness parameter *ω*, which regulates viral transcription and the transition of latent infected cells into a deeply latent state.

The steady states of the system depend on the effective reproduction number

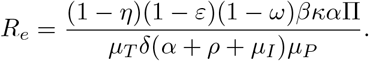

From this expression, it is evident that *R*_*e*_ decreases as the Tat inhibitor effectiveness *ω* increases. The threshold condition

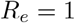

determines a critical value *ω*_*c*_ such that

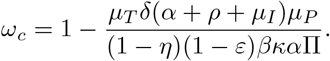

When *ω* < *ω*_*c*_, the effective reproduction number satisfies *R*_*e*_ > 1 and the infection persists within the host. When *ω* > *ω*_*c*_, the reproduction number becomes *R*_*e*_ < 1 and the virus-free equilibrium becomes globally stable.

The system therefore undergoes a transcritical bifurcation at *ω* = *ω*_*c*_, where stability shifts from the endemic equilibrium to the virus-free equilibrium.

The reproduction number *R*_*e*_ acts as a threshold parameter separating viral clearance from persistent infection. At *R*_*e*_ = 1, the virus-free equilibrium loses stability and an endemic equilibrium emerges. Fig 5 illustrates the forward transcritical bifurcation at the critical threshold *R*_*e*_ = 1, where the virus-free equilibrium loses stability and the endemic equilibrium emerges.

**Fig 5.**
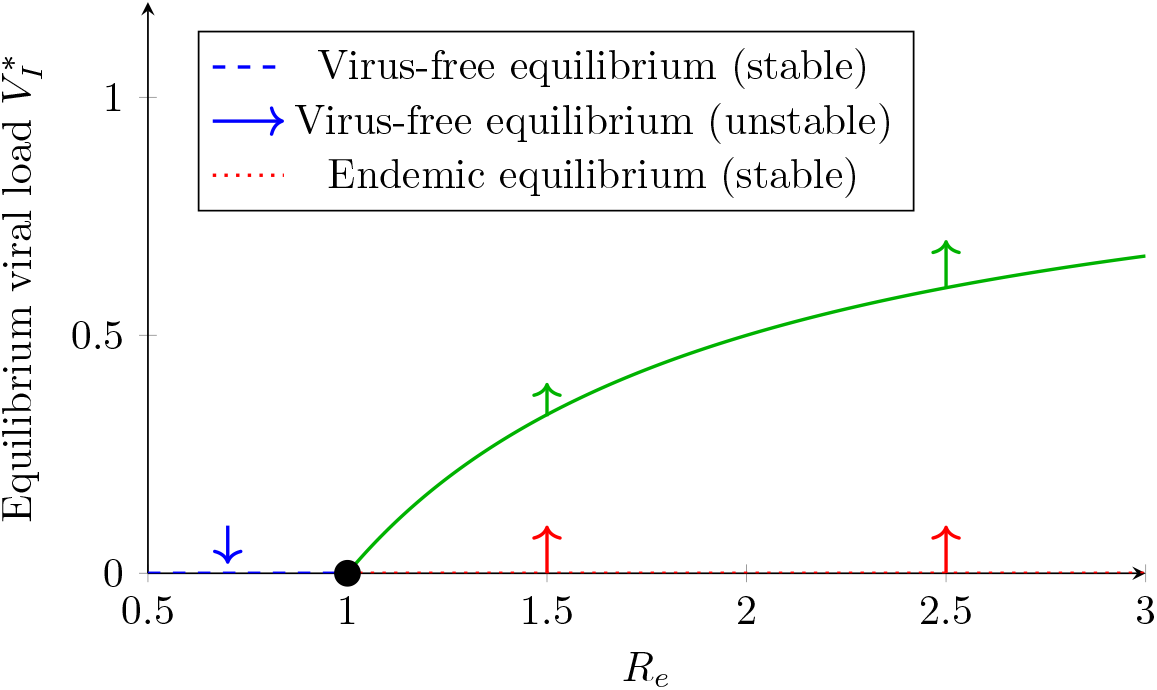
Forward transcritical bifurcation at *R*_*e*_ = 1. The virus-free equilibrium loses stability as *R*_*e*_ crosses unity, giving rise to a stable endemic equilibrium. Arrows indicate stability directions.

The graph implies that, the effective reproduction number *R*_*e*_ derived from system (1) determines the qualitative dynamics of the model. When *R*_*e*_ < 1, the virus-free equilibrium is locally asymptotically stable; when *R*_*e*_ > 1, a unique endemic equilibrium emerges through a forward transcritical bifurcation.

The system exhibits a forward bifurcation at *R*_*e*_ = 1 which provides a useful framework for interpreting the effects of different therapeutic interventions on HIV dynamics.

#### 3.9.2 Therapy-Dependent Interpretation of the Bifurcation

Fig 6 shows how therapies reshape the endemic equilibrium branch, even when *R*_*e*_ > 1. To further understand the impact of therapeutic interventions on HIV dynamics, we perform a bifurcation analysis of the model with respect to key parameters influencing viral replication and latency formation. In particular, we focus on the Tat inhibitor effectiveness parameter *ω*, which regulates viral transcription and the transition of latent infected cells into a deeply latent state.

**Fig 6.**
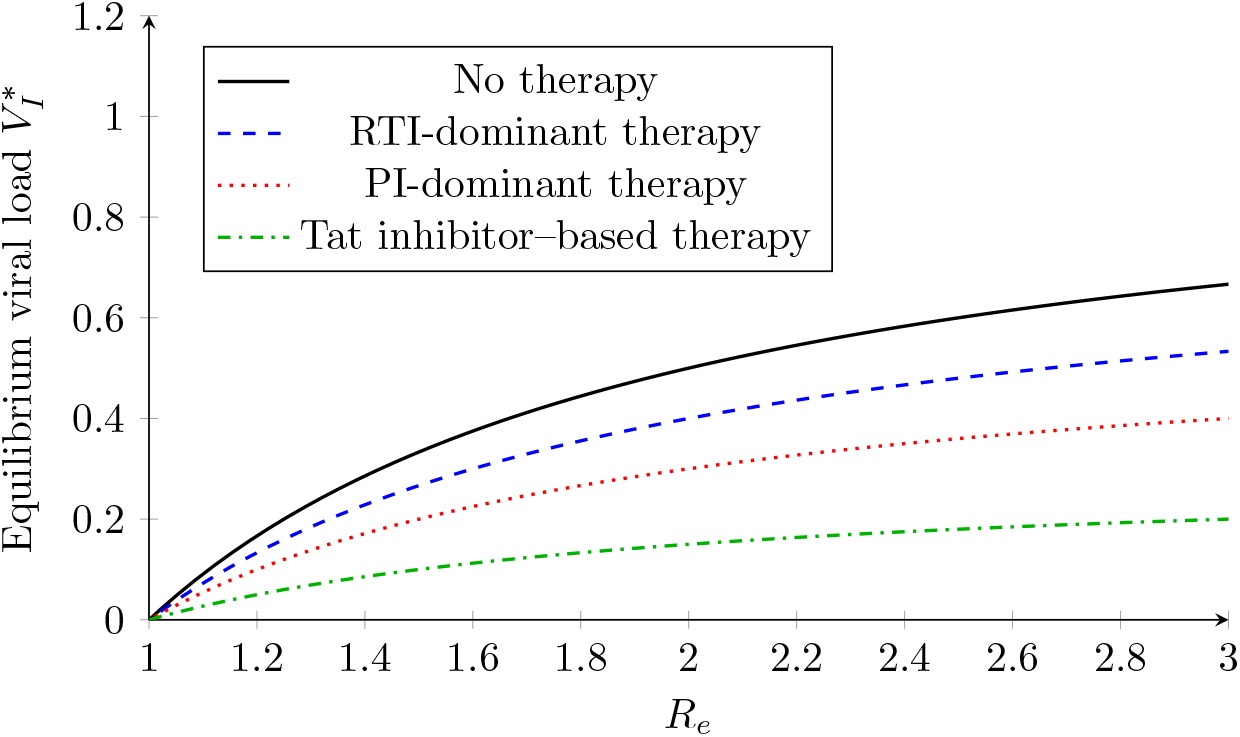
Therapy-dependent endemic equilibrium branches. Tat inhibition produces a pronounced reduction in the endemic viral set-point even when *R*_*e*_ > 1, while RTI-and PI-dominant therapies primarily scale viral load without altering the bifurcation structure.

Fig 7 illustrates the effect of Tat-inhibitor efficacy *ω* on the equilibrium viral load 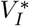 for a fixed *R*_0_ = 2.

**Fig 7.**
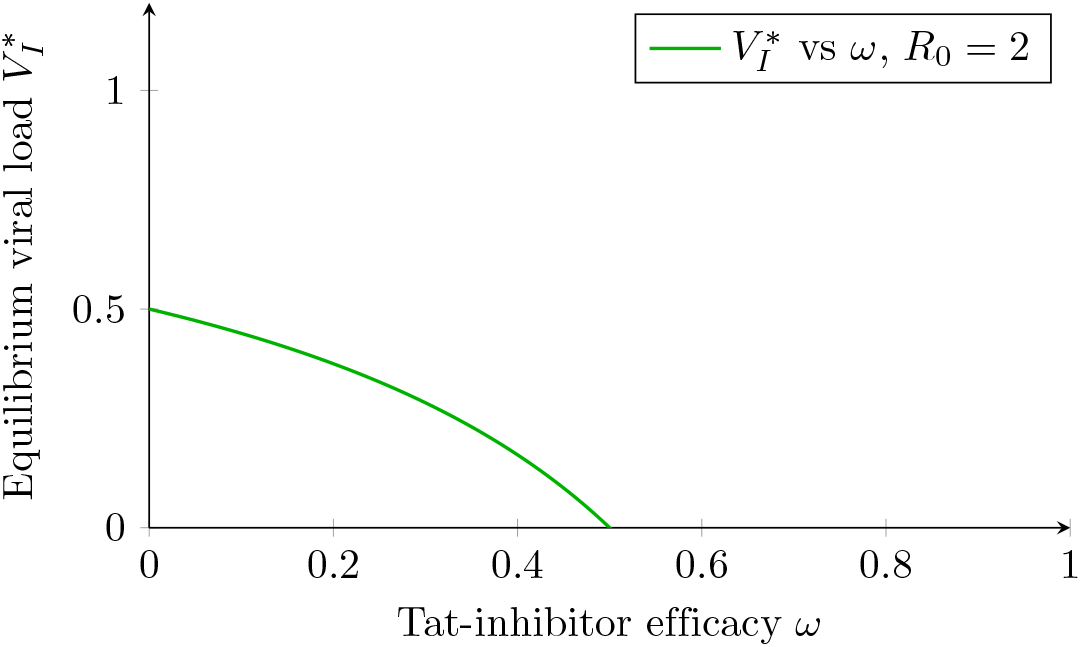
Effect of Tat-inhibitor efficacy *ω* on equilibrium viral load. Increasing *ω* reduces productive infection in system (1), thereby lowering the effective reproduction number and suppressing the equilibrium infectious virus level 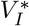 and potentially suppressing viral persistence.

#### 3.9.3 Interpretation of the Tat-inhibitor dependent Bifurcation Dynamics

The bifurcation diagram illustrates how the equilibrium viral load changes as the effectiveness of the Tat inhibitor increases. When the Tat inhibitor effectiveness *ω* is small, viral transcription remains active and productively infected cells continue to generate infectious virions. In this graph, the effective reproduction number exceeds unity (*R*_*e*_ > 1), and the infection persists within the host.

As *ω* increases, the transcription of viral RNA is increasingly suppressed. This reduces the transition of infected cells into productive infection and promotes the accumulation of deeply latent cells. Consequently, viral production declines and the equilibrium viral load decreases.

At the critical threshold *ω*_*c*_, the system undergoes a transcritical bifurcation where the endemic equilibrium collides with the virus-free equilibrium. Beyond this threshold (*ω* > *ω*_*c*_), the virus-free equilibrium becomes stable and the viral load approaches zero. Biologically, this result indicates that Tat inhibition shifts the infection dynamics from active replication toward transcriptional silence, reducing the contribution of the latent reservoir to viral rebound. In combination with other antiretroviral therapies, this mechanism may contribute to long-term viral control and reduction of the active viral reservoir.

## 4 Numerical Simulations

This section is aimed at numerically investigating the behavior of each compartment. The initial values of the model were set as *T*(0) = 1000, *I*(0) = 10, *I*_*L*_(0) = 5, *I*_*P*_ (0) = 5, *I*_*D*_(0) = 5, *V*_*I*_ (0) = 100 and *V*_*U*_ (0) = 0. The values of the parameters used were obtained from literature sources as shown in Table 3 below.

To complement the analytical results, numerical simulations were performed to investigate how therapeutic interventions influence viral replication, latent reservoir formation, and viral rebound dynamics. The simulations focus on the interaction between reverse transcriptase inhibitors, protease inhibitors, and Tat transcription inhibitors, with particular attention to their combined ability to reduce the effective reproduction number and suppress viral persistence. These visualizations provide insight into how different therapy combinations influence viral replication, latent reservoir formation, and viral eradication conditions.

### 4.1 Baseline Dynamics of Deep Latency (*I*_*D*_), Productive Infection (*I*_*P*_), and Viral Load (*V*_*I*_)

The comparative simulations under no treatment, conventional ART, and ART combined with Tat inhibition reveal distinct dynamical patterns in the deeply latent infected cells (*I*_*D*_), productively infected cells (*I*_*P*_), and infectious viral particles (*V*_*I*_). These trajectories illustrate how different therapeutic mechanisms influence both viral replication and the structure of the latent reservoir.

#### Dynamics without treatment

In the absence of therapy as depicted by the black line in Figure 8, the model predicts a rapid increase in productively infected cells (*I*_*P*_), which drives a corresponding exponential rise in infectious viral particles (*V*_*I*_). As viral replication proceeds, the latent infected cells reactivate and become productively infected, leading to gradual production of virus (*V*_*I*_). Over time, the system approaches an endemic equilibrium characterized by high viral load and sustained levels of productive infection.

**Fig 8.**
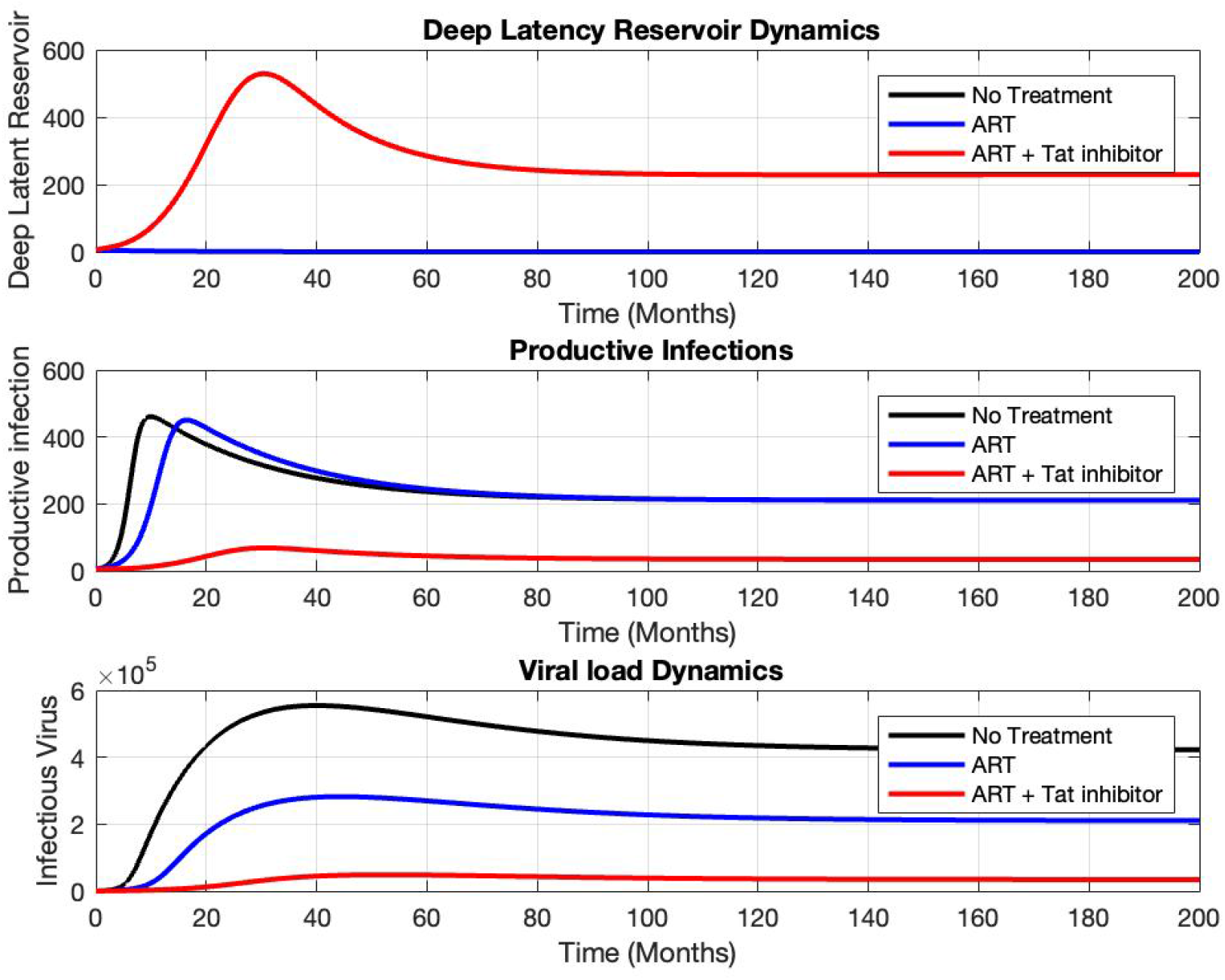
Evaluating the Efficacy of Therapies for Viral Load Suppression and Latency.

#### Dynamics under conventional ART

When antiretroviral therapy is introduced as shown by the blue line in Figure 8, the trajectories change slightly. The number of productively infected cells (*I*_*P*_) declines slightly due to the suppression of new infections but viral reactivation still persists . Consequently, the viral load (*V*_*I*_) decreases sharply and stabilizes at very low levels. However, the deeply latent compartment (*I*_*D*_) remains relatively stable at low levels over time. This behavior reflects the well-established limitation of conventional ART: while it effectively blocks viral replication, it does not eliminate latently infected cells which reactivate to productive infection.

#### Dynamics under ART combined with Tat inhibition

The introduction of Tat transcription inhibition alongside ART as shown by the red line in Figure 8, produces a distinct dynamical pattern. The number of productively infected cells (*I*_*P*_) declines rapidly and viral load (*V*_*I*_) falls to very low levels. However, the dynamics of the deeply latent compartment (*I*_*D*_) differ significantly. The simulations show a progressive increase in the deeply latent population accompanied by a stronger suppression of productive infection. This shift occurs because Tat inhibition suppresses viral transcription, reducing the likelihood that latently infected cells transition into productive infection. In general, the combined therapy leads to both reduced viral load and diminished reactivation potential of the latent reservoir.

Biologically, this mechanism corresponds to stabilization of transcriptionally silent proviruses, consistent with the proposed “block-and-lock” strategy in HIV cure research. By preventing transcriptional activation of integrated proviral DNA, Tat inhibition reduces the production of infectious virions and limits replenishment of the actively replicating viral pool.

Overall, the simulations suggest that effective HIV cure strategies may require coordinated interventions targeting all three processes simultaneously, which may potentially reduce the likelihood of viral rebound following treatment interruption.

### 4.2 Impact of Therapy Efficacies on Viral Rebound and Deep Latency Reservoir

Figure 9 illustrates the combined effects of therapy efficacies on viral rebound and the deep latent reservoir following treatment interruption. The results highlight how variations in infection inhibition (*η*), viral production inhibition (*ε*), and transcriptional inhibition (*ω*) jointly determine both the magnitude of viral resurgence and the structure of the latent reservoir. Heat maps provide a complementary visualization of therapy parameter combinations that minimize viral rebound and maximize on deep latency.

**Fig 9.**
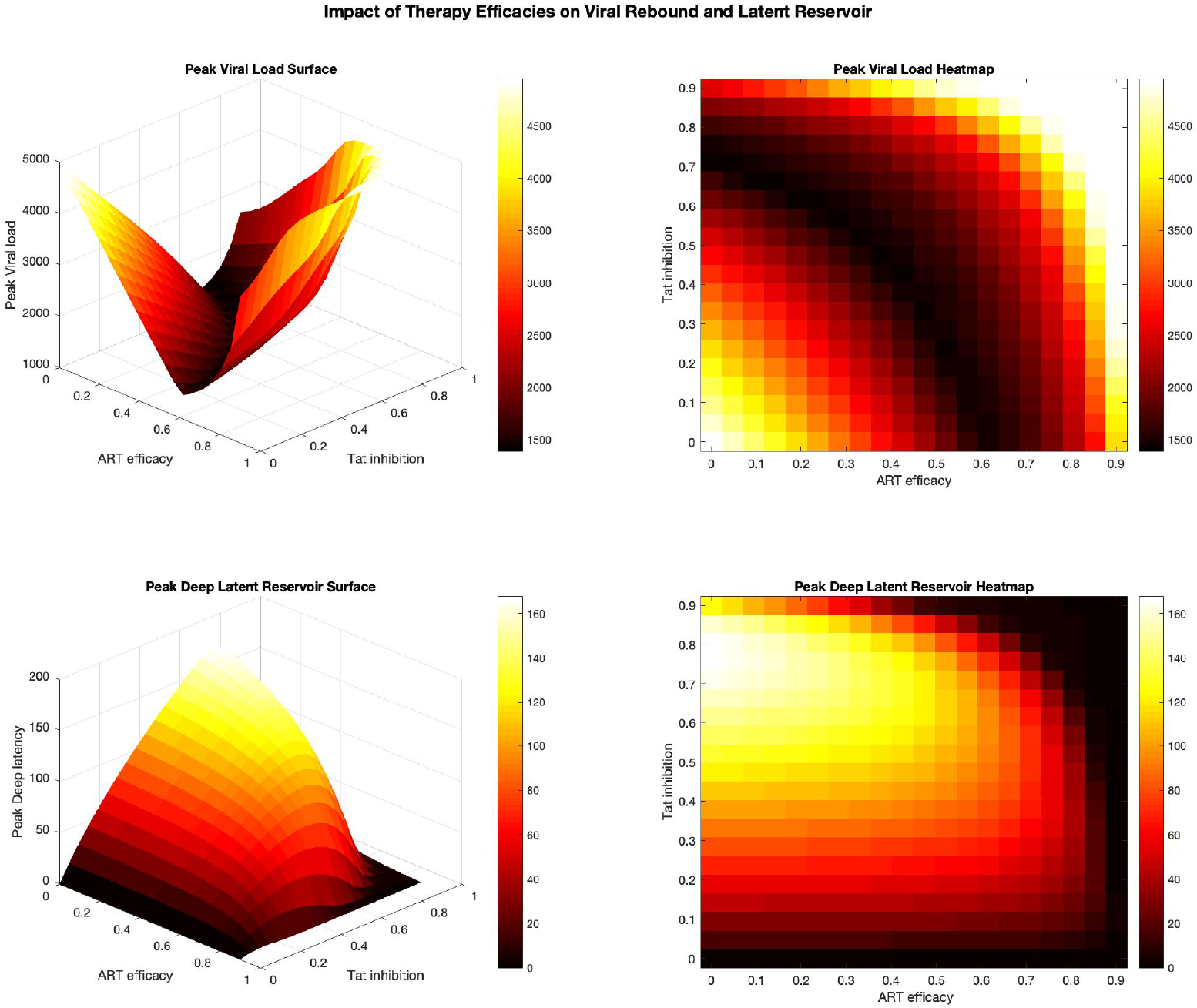
Impact of Therapy efficacies on viral Rebound and Deep latency reservoir.

#### Viral Rebound Dynamics

he viral load surface shows how combination of therapy, that is Tat inhibitor *ω* and ART minimize viral rebound. The surface shows that peak viral load after treatment interruption is strongly dependent on the combined efficacy of therapeutic interventions. When therapy efficacies are low, the model predicts a rapid and pronounced rebound in infectious viral particles (*V*_*I*_), reflecting the reactivation of latent reservoirs and the rapid expansion of productively infected cells.

As the efficacy of infection inhibition (*η*) and viral production inhibition (*ε*) increases, the magnitude of viral rebound decreases. This reduction occurs because fewer new infections are established and the production of infectious virions is suppressed. However, even at moderate levels of ART efficacy, viral rebound remains significant, indicating that conventional therapies alone may not sufficiently prevent reactivation.

In contrast, increasing transcriptional inhibition (*ω*) produces a more substantial reduction in rebound magnitude. The simulations show that high values of *ω* significantly flatten the rebound surface, indicating delayed and reduced viral resurgence. This effect arises because Tat inhibition suppresses transcriptional activation of proviral DNA, thereby limiting the transition of latent cells into productively infected states.

#### Deep Latent Reservoir Dynamics

The Deep latency surface shows effect of therapy parameters on maximal deep latency reservoir size. As transcriptional inhibition increases, the size of the deep latent reservoir also increases. This reflects a shift in the fate of infected cells: rather than progressing to productive infection, a larger fraction of cells enter or remain in a transcriptionally silent state.

While infection and production inhibition primarily reduce viral replication, they have a comparatively limited effect on the formation of deep latency. In contrast, transcriptional inhibition directly alters intracellular viral dynamics, promoting the stabilization of proviral genomes in a deeply latent state.

#### Implications for HIV Cure Strategies

These findings provide important insights for the design of HIV cure strategies. The results suggest that optimal therapeutic approaches may require a balance between suppressing viral rebound and managing the latent reservoir. In particular, combining transcriptional inhibition with conventional ART may reduce viral rebound to minimal levels while stabilizing the reservoir in a deeply latent state. This behavior aligns with the *block-and-lock* paradigm, in which viral transcription is permanently silenced to prevent reactivation.

These results highlight that effective HIV control requires not only suppression of viral replication but also strategic modulation of latency, emphasizing the need for therapies that simultaneously limit viral rebound while managing the long-lived reservoir.

### 4.3 Viral Set-Point Surface Under Combined Therapy Effects

To further investigate pairwise interactions between therapeutic mechanisms, parameter planes were constructed for combinations of the three therapy efficacies (*η, ε, ω*).

These planes illustrate how simultaneous modulation of infection inhibition, viral production inhibition, and transcription inhibition influences both viral load and latent reservoir dynamics.

The parameter planes for the deeply latent reservoir *I*_*D*_ represented by Figure 10a indicate that strong Tat inhibition plays a central role in driving infected cells toward deep latency. As a result, the deep latent reservoir grows while infectious viral production decreases. This result suggests that strong therapy may convert productive infection into deep latency. In contrast, RTIs and PIs primarily suppress viral replication without directly influencing the transition of latent cells into deeper transcriptionally silent states.

**Fig 10.**
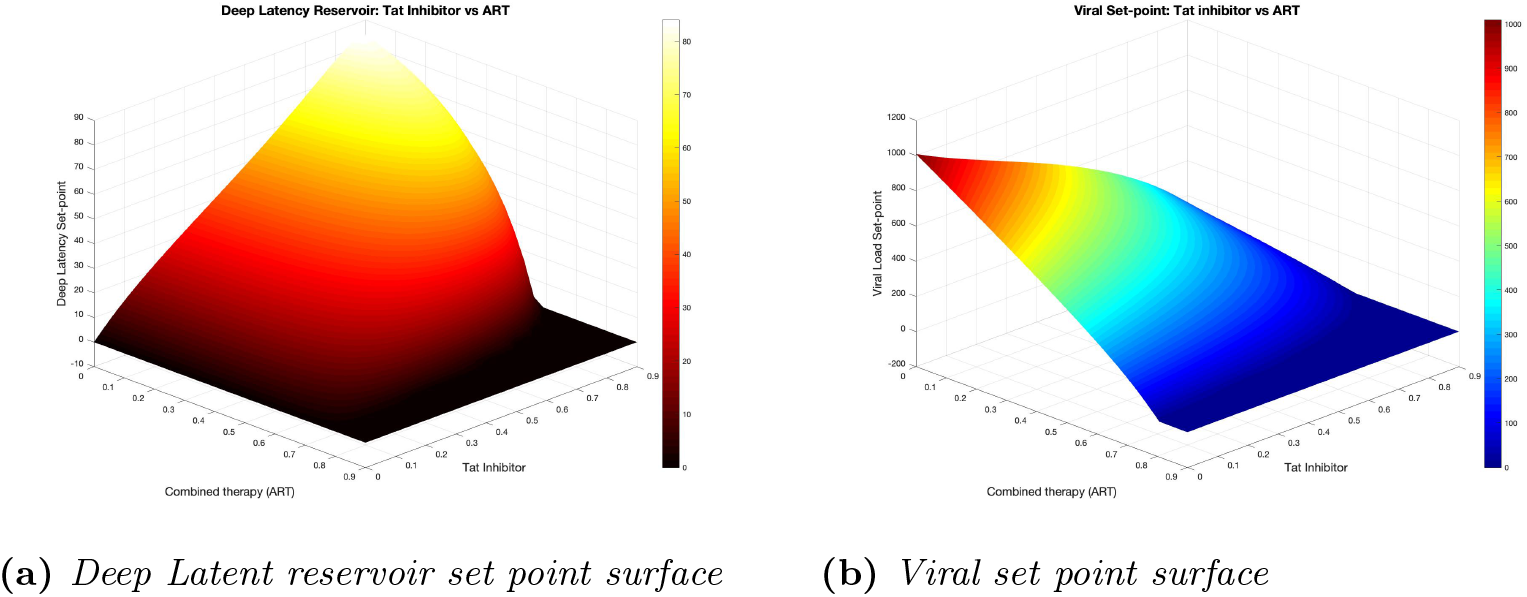
Viral Set-Point Surface Under Combined Therapy Effects.

Figure 10b shows the viral set-point surface when both infection inhibition and viral production inhibition act simultaneously in the presence of transcriptional suppression. The surface demonstrates a pronounced nonlinear decline in viral load as therapy efficacies increase.

In particular, once therapy efficacies exceed moderate thresholds, the viral set-point rapidly approaches very low levels. This sharp transition suggests that combination therapies targeting multiple stages of the viral life cycle are significantly more effective than therapies acting in isolation.

These findings align with the biological rationale of the block-and-lock strategy, in which transcriptional inhibitors are used to permanently silence proviral expression and reduce the likelihood of viral rebound.

Figure 11a illustrates the surface describing how the latent reservoir population varies with therapy efficacies. The simulations indicate that increasing transcription inhibition reduces the size of the actively replicating viral population but may initially increase the latent reservoir.

**Fig 11.**
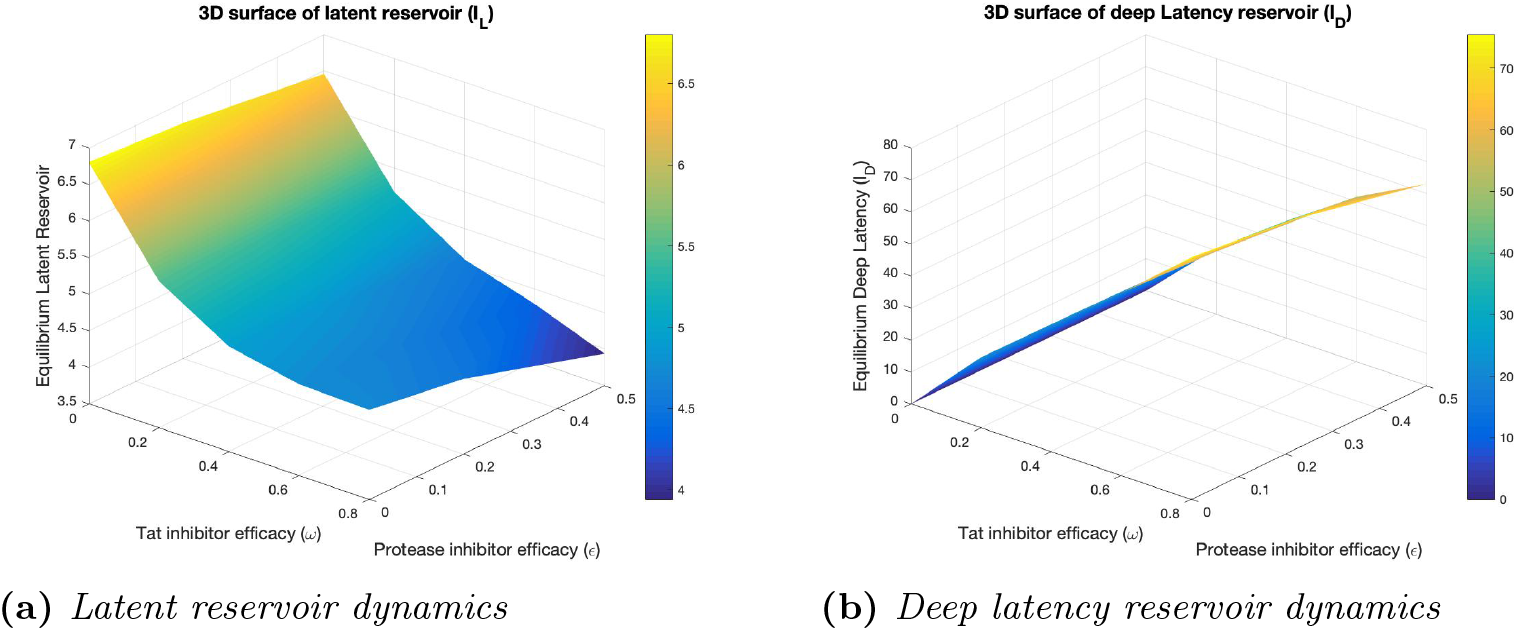
Effect of therapies on latent reservoirs.

This occurs because transcriptional suppression prevents productive infection, allowing infected cells to transition into a transcriptionally silent latent state. Such behavior is consistent with biological observations that strong transcriptional inhibition can stabilize latent reservoirs while suppressing viral replication.

Figure 11b presents the deep latent reservoir surface as a function of therapy efficacies. The surface demonstrates that increasing transcription inhibition promotes the accumulation of deeply latent infected cells.

This transition toward deeper latency reflects the mechanism underlying the *block-and-lock* therapeutic strategy, in which viral transcription is permanently suppressed to prevent reactivation of latent reservoirs. The results therefore suggest that strong Tat inhibition may contribute to long-term viral silencing even if complete eradication of infected cells is not achieved.

Tat inhibitors may lock latent virus into deep latency states. This is central to block-and-lock HIV cure strategies.

### 4.4 Eradication Boundary Surface

To identify therapy combinations capable of suppressing viral replication, a three-dimensional eradication boundary surface was constructed using the condition *R*_*e*_ = 1.

This surface separates parameter regions where viral persistence occurs (*R*_*e*_ > 1) from regions where viral replication cannot be sustained (*R*_*e*_ < 1).

The results show that individual therapies are generally insufficient to drive the system below the eradication threshold. Instead, effective viral control requires simultaneous action of multiple therapeutic mechanisms.

Figure 12 shows the three-dimensional eradication boundary surface defined by the condition *R*_*e*_ = 1. This surface separates the parameter region where infection persists (*R*_*e*_ > 1) from the region where viral replication cannot be sustained (*R*_*e*_ < 1).

**Fig 12.**
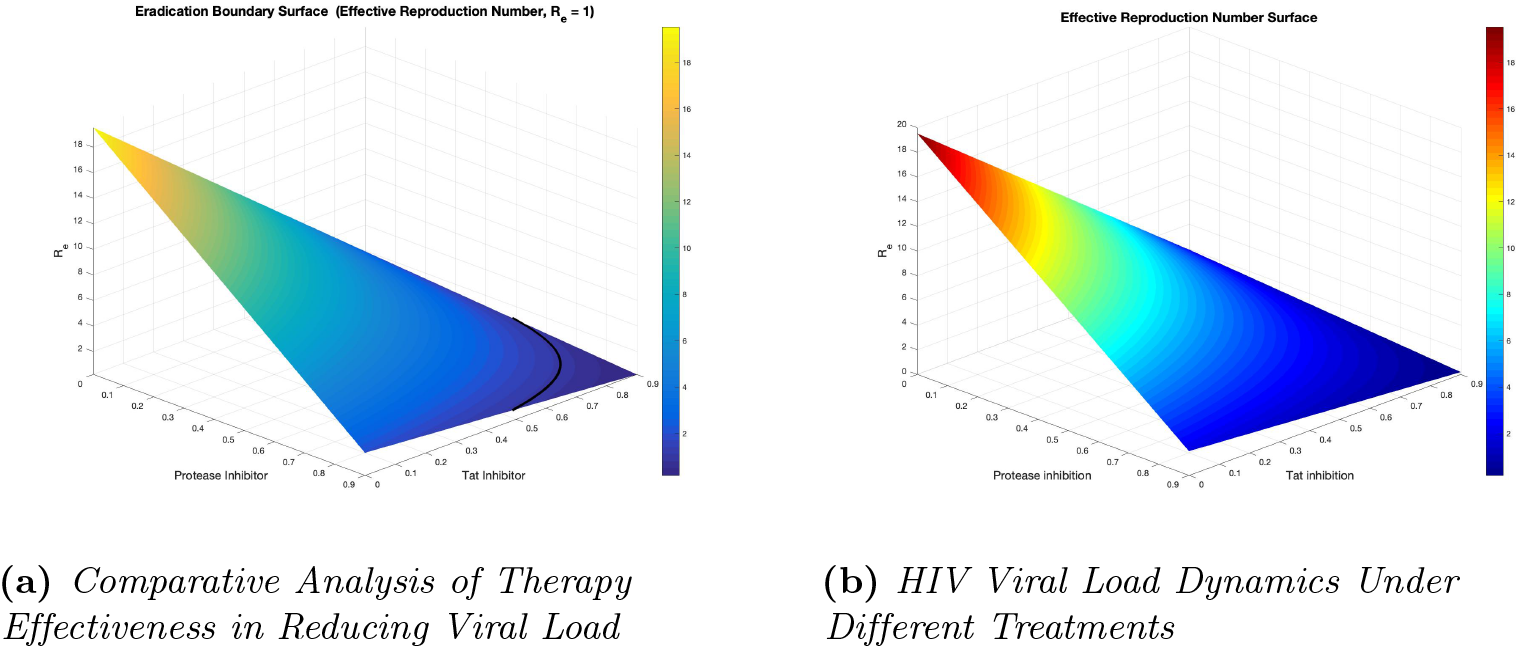
HIV Eradication Boundary Surface.

Points above the surface correspond to therapy combinations that reduce the effective reproduction number below unity, resulting in viral decline toward the virus-free equilibrium. The surface therefore provides a quantitative boundary identifying combinations of therapy efficacies required to achieve viral suppression.

These findings highlight the synergistic potential of Tat inhibitors in multi-drug HIV therapy and support ongoing efforts to integrate transcription-targeting drugs into future treatment strategies.

### 4.5 Three-Dimensional Therapy Cube

The therapy cube represented by Figure 13 presents a three-dimensional visualization of the therapy parameter space defined by infection inhibition (*η*), viral production inhibition (*ε*), and transcription inhibition (*ω*).

**Fig 13.**
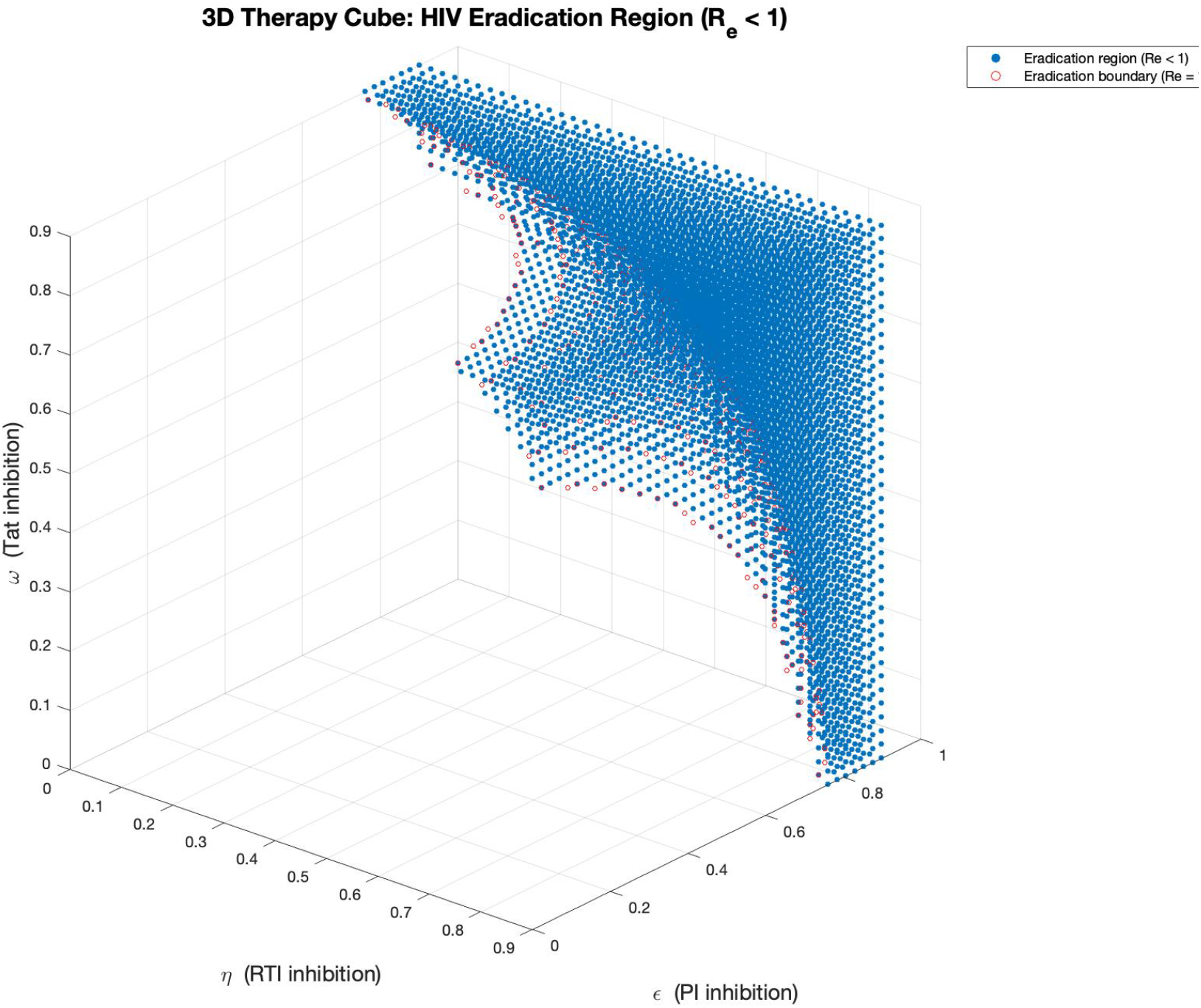
Therapy Cube Showing HIV Eradication Region.

Within this cube, the region satisfying *R*_*e*_ < 1 represents combinations of therapies capable of suppressing viral replication, that is, the viral eradication domain.

The therapy cube highlights that viral eradication occurs only when multiple therapeutic mechanisms act simultaneously. When any single therapy efficacy is low, the system remains in the persistent infection region. However, when infection inhibition, viral production inhibition, and transcription inhibition are all sufficiently strong, the system enters the eradication region where viral replication cannot be maintained. The cube demonstrates that eradication is unlikely when only a single therapy mechanism is applied [29, 30, 31, 32]. Instead, successful elimination typically occurs when multiple therapies act together, each contributing to the reduction of the effective reproduction number [33, 34, 35]. This visualization highlights the importance of combination therapy, which reduces viral replication through multiple independent mechanisms.

## 5 Discussion

This study presents a mathematical framework for investigating the *in vivo* dynamics of HIV infection under multiple therapeutic mechanisms, with particular emphasis on the role of Tat inhibition in regulating viral transcription and latency formation. The analytical and numerical results illustrate how different therapeutic mechanisms interact to influence viral replication, reservoir dynamics, and long-term infection outcomes. The derived effective reproduction number *R*_*e*_ provides a fundamental threshold that determines whether infection persists or declines. When *R*_*e*_ > 1, viral replication remains self-sustaining and infection persists within the host. In contrast, when *R*_*e*_ < 1, viral replication cannot be maintained and the infection gradually declines toward the virus-free equilibrium. This threshold provides an important quantitative framework for evaluating the effectiveness of therapeutic interventions in suppressing HIV replication. The three-dimensional therapy surfaces and heat maps introduced in this study provide deeper insight into how different drug mechanisms interact to regulate viral dynamics. These visualizations demonstrate that increasing the efficacy of infection inhibition (*η*), viral production inhibition (*ε*), or transcriptional inhibition (*ω*) reduces the effective reproduction number and suppresses viral replication. However, the simulations clearly indicate that strong viral suppression is most efficiently achieved when multiple therapies act simultaneously, highlighting the importance of combination therapy strategies.

The viral set-point surfaces reveal how combinations of therapeutic parameters influence equilibrium viral load. Increasing Tat inhibitor efficacy significantly lowers the viral set-point, particularly when combined with strong viral production inhibition. These results support experimental observations suggesting that targeting transcriptional activation can substantially reduce viral replication by limiting the generation of productively infected cells.

The latent and deep latent reservoir planes provide additional biological insight into the mechanisms underlying viral persistence. The simulations indicate that strong Tat inhibition not only suppresses viral production but also promotes the transition of infected cells into a deeply latent transcriptionally silent state. This behavior aligns with the emerging *block-and-lock* strategy, which seeks to permanently silence HIV transcription rather than eliminate all infected cells.

The eradication boundary surface and therapy cube provide a comprehensive visualization of the parameter combinations required to achieve *R*_*e*_ < 1. These figures demonstrate that eradication regions occupy only a limited portion of the therapy parameter space when individual drug mechanisms act alone. However, when infection inhibition, viral production inhibition, and transcriptional inhibition act together, the eradication region expands substantially. This finding emphasizes that multidrug therapeutic strategies targeting different stages of the viral life cycle are essential for achieving durable viral suppression.

Another important finding of this study relates to viral rebound following treatment interruption. The simulations confirm that interruption of therapy leads to rapid viral rebound driven by the reactivation of latent reservoirs. However, treatment regimens incorporating transcriptional inhibition significantly delay viral rebound and reduce rebound viral peaks. These results suggest that Tat-based therapies may enhance long-term viral control and reduce the risk of viral resurgence after therapy interruption. Taken together, the findings demonstrate that targeting viral transcription provides an important complementary mechanism to existing ART strategies. While traditional ART primarily blocks viral entry and maturation processes, Tat inhibitors suppress transcriptional activation and promote deeper latency states. Integrating these mechanisms within combination therapy may simultaneously reduce viral replication, suppress viral rebound, and limit the contribution of latent reservoirs to ongoing infection.

Overall, this study highlights the potential of combining transcriptional inhibition with conventional antiretroviral therapy as a promising strategy for suppressing viral replication and promoting durable viral control. The results suggest that Tat inhibitor-based combination therapies may contribute significantly to emerging functional cure strategies for HIV, particularly within the framework of block-and-lock approaches aimed at permanently silencing viral transcription.

While the present model provides important insights into the interaction between transcriptional inhibition and conventional antiretroviral therapies, several extensions may further improve its biological realism and predictive capability. First, future work could incorporate explicit immune responses, particularly cytotoxic T lymphocyte (CTL) dynamics, which play an important role in controlling infected cells and shaping viral rebound dynamics. Second, the current model assumes constant therapy efficacies; incorporating pharmacokinetic and pharmacodynamic processes would allow a more realistic representation of drug concentrations and treatment adherence patterns. Third, stochastic effects associated with the activation of latent reservoirs could be included to better capture the variability observed in viral rebound following treatment interruption. In addition, incorporating spatial heterogeneity and tissue-specific viral reservoirs, such as lymphoid tissues and the central nervous system, would provide a more comprehensive description of HIV persistence within the host. Finally, future studies may explore optimal therapy scheduling and adaptive treatment strategies aimed at minimizing viral rebound while promoting deeper latency states. Such extensions would help bridge the gap between theoretical modeling and clinical applications, providing further insight into the design of effective HIV cure strategies.

### 5.1 Implications for Current HIV Cure Strategies

The results obtained in this study have important implications for emerging HIV cure strategies. In particular, the analysis highlights the critical role of transcriptional inhibition in complementing conventional antiretroviral therapy. While standard ART regimens primarily target viral entry, reverse transcription, and virion maturation, they do not directly silence viral transcription within infected cells. The simulations presented here show that incorporating Tat inhibition can substantially reduce viral set-points, delay viral rebound following treatment interruption, and promote the transition of infected cells toward deeper latent states. These findings align closely with the *block-and-lock* strategy, which aims to permanently silence HIV transcription and prevent viral reactivation from latent reservoirs. In contrast to eradication approaches such as *shock-and-kill*, which attempt to activate and eliminate latent infection, the present results suggest that reinforcing transcriptional silencing may provide a viable pathway toward durable viral control. The therapy surfaces and eradication cube further demonstrate that achieving sustained viral suppression requires coordinated targeting of multiple stages of the viral life cycle, including infection, viral production, and transcriptional activation. Together, these insights support the development of multidrug therapeutic strategies that integrate conventional ART with transcription-targeting agents, offering a promising direction for advancing functional cure approaches for HIV infection.

## Acknowledgments

The authors acknowledge with gratitude the support from Strathmore University in the production of this manuscript.

